# The GA4GH Categorical Variation Representation Specification: A Unified Computational Framework for Reasoning over Genomic Variant Categories

**DOI:** 10.64898/2026.02.10.705161

**Authors:** Daniel Puthawala, Brendan Reardon, Lawrence Babb, Kori Kuzma, James S. Stevenson, Wesley A. Goar, Robert H. Dolin, Robert R. Freimuth, Catherine Procknow, Beth Pitel, Parijat Kundu, Akash Rampersad, Eliezer Van Allen, Alex H. Wagner

**Affiliations:** The Steve and Cindy Rasmussen Institute for Genomic Medicine, Nationwide Children’s Hospital, Columbus, OH; Department of Medical Oncology, Dana-Farber Cancer Institute, Boston, MA; Cancer Program, Broad Institute of MIT & Harvard, Cambridge, MA; Medical and Population Genetics, Broad Institute of MIT and Harvard, Cambridge, MA; Elimu Informatics, El Cerrito, CA; Mayo Clinic, Rochester, MN; Epic Systems,Verona, WI; Velsera, Pune, India; OakBioinformatics LLC, Fairfax, VA; Division of Medical Sciences, Harvard University, Boston, MA; Departments of Pediatrics and Biomedical Informatics, Ohio State University College of Medicine, Columbus, OH

## Abstract

Categorical variants, or sets of genomic alterations constrained by shared properties, are pervasive across clinical, regulatory, and research domains in the biomedical ecosystem, yet their inconsistent and non-computable representation hinders data interoperability and clinical interpretation. We surveyed genomic knowledgebases spanning regulatory approvals and the biomedical literature and found that categorical variants underpin a substantial proportion of clinical genomics knowledge, but are largely described using incompatible bespoke models. To address this, we developed the GA4GH Categorical Variation Representation Specification (Cat-VRS), a constraint-based framework that provides a unified computable representation for both precise and intentionally broad categories across molecular and systemic variant domains. Cat-VRS enables harmonization of genomic knowledgebases, computable category-based search, and automated matching between assayed variants and categorical entities in clinical and research contexts. By providing a principled, extensible model for categorical variation, Cat-VRS enables computable reasoning over genomic variant categories and establishes a foundation for the standardized representation and exchange of genomic knowledge.

## Introduction

The promise of genomic medicine depends on our ability to recognize assayed genomic alterations as belonging to clinically significant variant classes. Yet linking observed variants to these shared knowledge representations remains challenging. Unstructured data, divergent nomenclatures, and ambiguous conceptual descriptions all fragment how variants are described, impeding automated comparison and interpretation. Bridging these representations is essential for efficient, accurate, and equitable delivery of genomic testing and precision medicine to serve patients.

Associating genomic variants with knowledge is complicated by multiple valid variant representations. A single alteration can be expressed in multiple equivalent ways depending on transcript choice, alignment conventions, or reference assembly, thereby creating a many-to-many mapping between observed data and existing knowledge representations. This was alleviated for sequence variants through the introduction of canonical variant identifiers from which alternative representations can be expressed relative to^1,2^. This notion of grouping related assayed variants into a larger class can itself be generalized to the idea of **categorical variants:** sets of related variants that share functional, structural, and/or clinical properties^3,4^. For example, there are several known variants constituting *EGFR* exon 19 deletions^5^, and collectively they represent examples of a therapeutically actionable category associated with sensitivity to several EGFR inhibitors in non-small cell lung cancer (Fig. 1). Categorical variants are currently described within regulatory approvals, clinical trial enrollment criteria, journal reports, pharmacogenomics resources, and functional studies, and report on diverse types of categorical variants such as *TPMT* deficiency^6^, *BRAF* Class 2 variants^7^, *SETBP1* exon 4 variants^8^, *CYP2D6* *5^9^, and *PTEN* loss of function^10^, respectively.

**Fig. 1:**
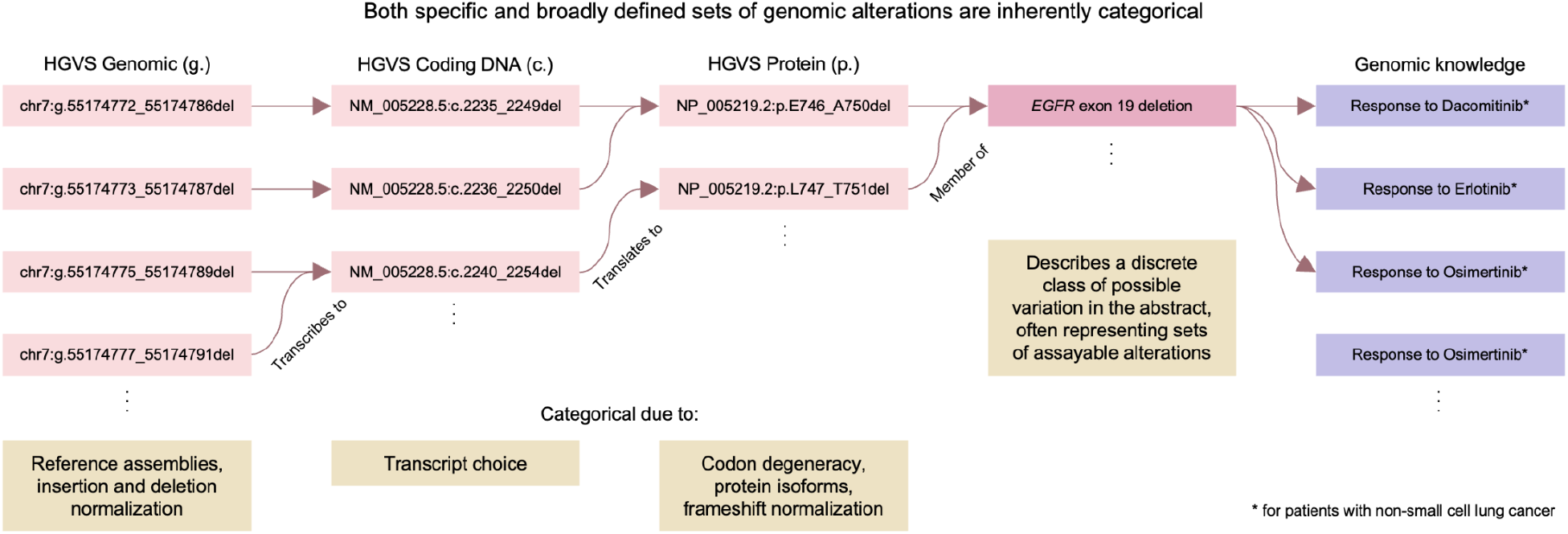
Both specific and broadly defined sets of genomic alterations are inherently categorical. Even a single, specific genomic alteration is inherently categorical because it can be validly expressed in multiple ways depending on molecular context and representational conventions. A sequence variant may appear differently across reference assemblies, normalization schemes, or transcript choices. Protein-level expressions vary with isoforms, codon degeneracy, and frameshift normalization, while distinct nomenclatures (such as HGVS or pharmacogenomic star alleles) provide additional layers of representation. Likewise, broadly defined sets of genomic alterations such as ‘*EGFR* exon 19 variants’ may encompass numerous or non-finite potential members, making them historically difficult to represent computably within genomic knowledgebases. Together, these observations illustrate that both specific and broad genomic alterations possess categorical structure.

While the Global Alliance for Genomics and Health (GA4GH) Variation Representation Specification^11^ (VRS) established a foundation for unambiguous representation of sequence-level variants, extending these principles to categorical variation has remained an unsolved problem. Feasibility studies of standards-based variant annotation of FHIR Genomics and early GA4GH Variant Annotation models highlighted the complexity of annotation increases dramatically for structural and systemic variant types, including copy number variants and rearrangements^12^. Any framework for categorical variants therefore must accommodate properties such as molecular function, ontology, or effect on biological systems, and must additionally accommodate both precise and intentionally vague representations.

To address this representational gap, we developed the **Categorical Variant Representation Specification (Cat-VRS; pronounced “cat-verse”)**, a new standard from the GA4GH Genomic Knowledge Standards (GKS) Work Stream. Cat-VRS provides a constraint-based framework for modeling both specific and broad categorical variants. In this model, each constraint defines a property that filters the set of possible genomic alterations matching a described categorical variant, enabling both precise and intentionally broad representations. Here, we analyze the landscape of genomic knowledgebases spanning regulatory approvals and the biomedical literature, formalize a typology of categorical variants, and describe how these insights informed the design and implementation of Cat-VRS.

## Results

### Landscape analysis of categorical variation

To characterize the prevalence and diversity of categorical variant representation, we surveyed knowledgebases, clinical resources, data standards, informatics tools, and registries that either store, reference, or operate on genomic alterations (Supplementary Table 1)^1–4,11,13–37^. We observed that categorical variants are not confined to any single domain. Both somatic and germline knowledgebases^3,15,16,18,20,38^, API-driven annotation platforms^13,14^, electronic health record infrastructures^39,40^, clinical trial registries^41^, and international data standards^2,31,42,43^ all rely on categorical variant definitions. Across these diverse systems, categorical variants serve as core organizational units for biomarker definition, clinical eligibility determination, and evidence interpretation. Yet, because no standardized or computable representation exists, many of these systems implement their own idiosyncratic category models; for example, CIViC’s variant type system^44^ and JaxCKB’s bespoke category schema^3^. As a result, semantics drift across resources, connections between categories and their members are frequently lost, and harmonization remains difficult. These observations motivated a deeper examination of how categorical variants appear in two contexts where they have strong clinical impact: regulatory approvals and genomic knowledgebases.

Oncology approvals from the US Food and Drug Administration were reviewed for those involving biomarkers as part of the indication (“precision oncology”), with special focus on approvals involving broad classes of variation rather than single genomic alterations. We observed that 83 of 236 precision oncology approvals (35.17%) referenced categorical variants as part of the approved indication (Fig. 2a; Methods). The specificity of these categories varied widely. Some approvals referenced specific structural or location-defined categories (for example, *EGFR* exon 19 deletions, *FGFR2* rearrangements), whereas others relied on functionally or clinically defined categories such as “susceptible *IDH1* variants.” In clinical practice, molecular pathologists, variant curation teams, and tumor boards must determine whether an observed variant satisfies these categorical definitions, often without a computable standard to assist them. An effort that continues to grow more resource intensive as both known clinically-relevant biomarkers and clinically-available therapies increase over time^45^.

**Fig. 2:**
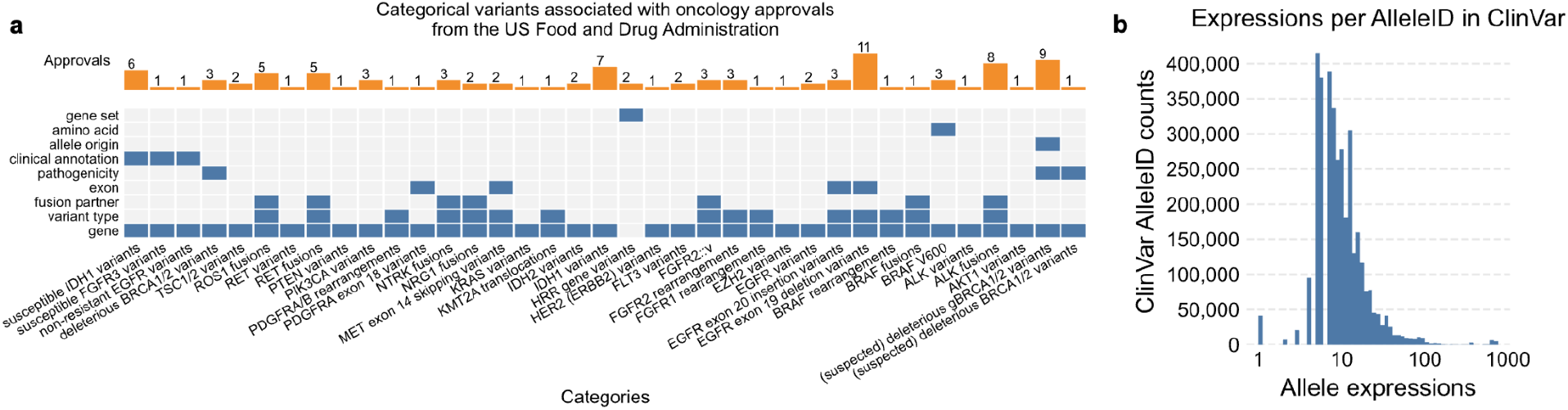
Prevalence of categorical variants associated with genomic knowledge. (a) 83 of 236 (35.17%) precision oncology approvals from the US Food and Drug Administration referenced broad categories of genomic variation. (b) Individual sequence variants are also inherently categorical due to expression multiplicity. While the median number of HGVS expressions per Allele in ClinVar was 9, 0.68% have more than 100 representation expressions.

Genomic knowledgebases curate statements involving categorical variants from both regulatory approvals and scientific literature. They do so by cataloging categories directly, by associating individual variants with categorical evidence statements, or a combination of the two, frequently losing explicit links between categories and their members in the process. For example, while precision oncology knowledgebases contain genomic knowledge statements for “*NTRK1/2/3* fusions” broadly^16,24,46^, COSMIC has curated 26 distinct fusions involving *NTRK1/2/3*^15^. Likewise, CIViC maintains 71 variant types based on Sequence Ontology terms^16,26^, which collectively encompass 2,439 curated variants, reflecting many of the actionable concepts used by expert curation panels.

Codon degeneracy, transcript variability, isoforms, and reference alignment differences all contribute to expression multiplicity of individual sequence variants. For example, NM_000518.5(HBB):c.20A>T(p.Glu7Val), a pathogenic variant for sickle cell anemia, has 11 distinct HGVS expressions displayed in ClinVar (8 Nucleotide, 3 protein) in addition to its MANE Select representation^17,21,22^. Indeed, while the median number of HGVS expressions per ClinVar Allele is 9, the distribution has a long tail: 24,221 of 3,568,536 (0.68%) having greater than 100 (Fig. 2b; Methods), due to factors such as multiple sequence contexts, transcript and protein isoforms, insertion and deletion alignment, and alternative reference assemblies.

While tools such as VariantValidator^47^ and the Variation Normalizer^28^ can mitigate these expression-level inconsistencies by harmonizing molecular representations, they cannot resolve the broader challenge: many categorical variants are not sequence-resolvable at all. Categories based on functional impact (for example, loss-of-function *PTEN* variants), structural configuration (for example, rearrangements such as large-scale sequence duplications, inversions, and translocations), biological consequence (for example, “activating” *BRAF* variants), or clinical criteria (for example, “susceptible” *IDH1* variants) do not correspond to a single molecular allele nor to a simple set of sequence changes. As a result, improving allele-level representation alone cannot capture the full space of categorical variation encountered in clinical genomics. A computable framework that unifies both precise alleles and non-sequence-resolvable categories is needed to provide a way to express shared properties while preserving intentional ambiguity.

### A formal analysis of categorical variation in the CIViC knowledgebase

A central challenge in developing a standardized representation for categorical variants is that existing knowledgebases often employ overlapping, inconsistently defined categories. To clarify the semantics of these categories, we performed a formal qualitative analysis of the variant types curated in the CIViC knowledgebase^16^. CIViC is publicly available, widely used, and integrates community-driven submissions with expert panel review, providing a unique window into both the data model and the collective intuitions of the field^16^. Our goal was to disentangle how these categories are defined, what properties they entail, and how they relate across biological levels of description.

We began by constructing an ontology log (olog)^48^, where each CIViC variant type was represented as a set object and the definitional relationships among them as set morphisms (Supplementary Fig. 1). This formalism identifies the minimal set of properties that govern category membership. For example, the distinction between in-frame and frameshift deletions reduces to a single constraint on the length of the deleted sequence, while both remain specializations of the broader class of deletions.

Because many categorical variants ultimately entail particular kinds of change, such as changes in sequence, location, frame, quantity, or function, we inverted the olog to generate a typology in which these axes of variation and the biological level of representation define a coordinate space. In this typology, the axes are defined by the kind of change and the biological level at which it occurs, forming a coordinate space in which CIViC’s variant types can be placed (Fig. 3). Categories that span multiple axes, such as gene fusions or structural inversions, occupy multiple positions and appear as connected nodes.

**Fig. 3:**
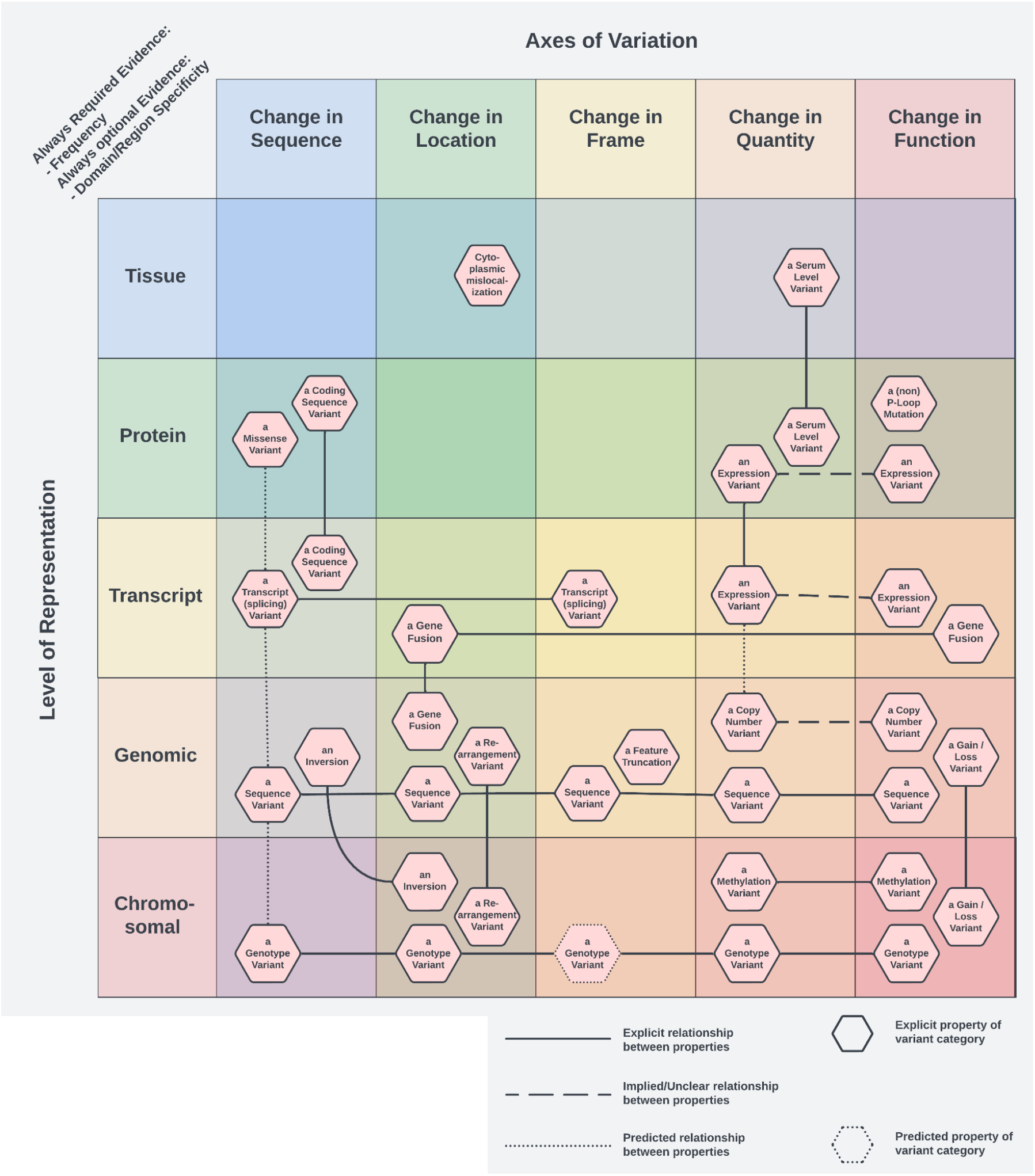
A typology of variant categories in the CIViC knowledgebase. A projection of CIViC’s categorical variant typology into a coordinate space defined by axes of variation (the kind of change) and the biological level at which that change is represented. By inverting the underlying olog, each variant category appears at all positions corresponding to its defining properties, with links indicating multi-axis membership. This layout makes the structure of CIViC’s categories visually intuitive, revealing where categories cluster redundantly, where conceptually plausible categories are absent, and where parallel patterns emerge across biological levels. The typology summarizes the same formal relationships captured in the olog but in a form that exposes representation gaps and shared semantics, providing the conceptual substrate for the constraint-based representation model implemented in Cat-VRS.

The resulting coordinate typology provides a coherent conceptual map of CIViC’s variant types. By making both the axes of change and the level of biological representation explicit, it exposes **categorical gaps** (cells where a plausible category exists conceptually but is absent in CIViC, such as protein-level rearrangement variants), clarifies **redundancy** (cases where categories differ only superficially, such as coding sequence variants versus transcript variants), and highlights **parallelism** (including analogous categories defined at different biological levels, such as methylation variants and copy-number variants). By grounding the formal analysis variants in CIViC in this way, we obtained a principled basis for modeling the semantics of categorical variants that is faithful both to their practice usage and their underlying biological structure. This analysis provided the conceptual foundation for the constraint-based representation that underpins Cat-VRS, enabling a systematic and extensible way to encode categorical variants across diverse knowledgebases.

### Design of a constraint-based representation model

Insights from the CIViC olog analysis highlighted that categorical variants can be defined by the minimal properties that determine category membership, rather than through reliance on the enumeration of member variants. This observation motivated a representation strategy that could capture both precise alleles and broad, intentionally vague classes within a single computable framework. To achieve this, we developed the Cat-VRS, a constraint-based data model within the suite of GA4GH genomic knowledge standards.

Historically, two representational paradigms have dominated genomic variant modeling. **Top-down typologies**, such as those used in CIViC or HGVS, organize variants into predefined categories (for example, deletions, fusions, copy-number changes). These systems are intuitive for curators but are also rigid, not easily extensible, and prone to creating artificial distinctions between closely related variant classes. In contrast, **bottom-up molecular models** like SPDI and allele-centric VCF encodings emphasize precise, deterministic representations of specific alterations, but provide limited support for categories that span multiple biological levels or capture functional or clinical concepts. Neither paradigm alone can represent the full spectrum of categorical variation encountered in genomic medicine.

Cat-VRS synthesizes these approaches by treating categorical variants as sets defined by composable constraints (Fig. 4). Each **constraint** represents a property that must be satisfied by any variant belonging to that category. For example, different constraints could require that all members relate to a specific allele, a region of a reference sequence, a copy-number change, a structural configuration, or a functional consequence (Table 1). Constraints can be combined to represent categories ranging from highly specific (for example, a single VRS Allele) to deliberately broad (such as, “variants affecting *PTEN* function” or “any rearrangement involving *FGFR2*”). Because constraints operate over shared properties rather than enumerated members, Cat-VRS supports both precision and ambiguity in a principled, computable way.

**Fig. 4:**
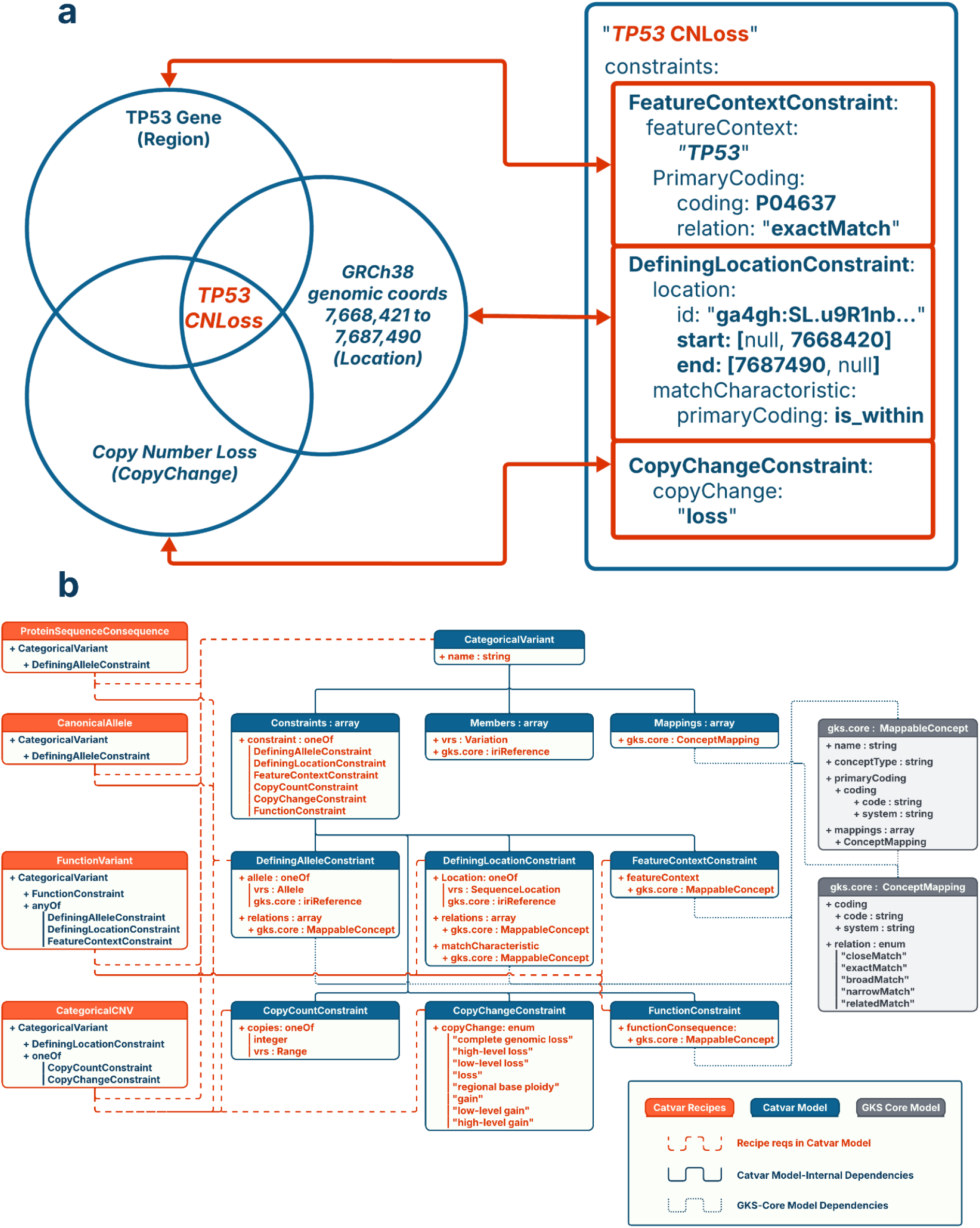
Constraint-based representation of a categorical variant in Cat-VRS. Cat-VRS models categorical variation as intensional sets through the intersection of their component properties via constraints, rather than enumeration of member alleles or reliance on fixed typologies. (a) The category “TP53 CNLoss” is represented by three modular constraints: *FeatureContext, DefiningLocation*, and *CopyChange*. Each constraint encodes a salient property of the category in a computable form. These constraints correspond intuitively to the biological features that characterize this variant class: being located within the *TP53* gene region, corresponding to a specific sequence location on GRCh38, and constituting a net loss of copy number. When composed within a single *CategoricalVariant* object, the constraints operate conjunctively, defining the set of variants that satisfy all required properties. (b) Cat-VRS currently supports 6 constraints (blue) and backwards compatibility with data classes from GA4GH VRS and GA4GH GKS-Core (gray), for interoperability with other GA4GH standards. Cat-VRS also supports Categorical Variant Types (orange), which allow Cat-VRS adopters to freely specify the properties required for certain types of *CategoricalVariant* objects for their local implementation. Some example types are included in the reference implementation, such as the *CanonicalAllele* and *CategoricalCNV* classes which stipulate the properties required of *CategoricalVariant* objects submitted to the ClinVar knowledgebase.

**Table 1:** Categorical Variant Representation Specification features and product roadmap. Cat-VRS represents categorical variants using modular constraints, each capturing a specific property in a computable form, including allele identity, genomic location, feature context, copy state, or functional effect. This table summarizes the constraint types currently released and under development along with their maturity level and a brief description of the property each encodes. It also lists current Categorical Variant Types, which provide reusable schemas for composing common forms of categorical variation from these underlying constraints for specific implementations. All described Data Classes are relative to the 1.0.0 Release on Cat-VRS’ GitHub repository.

| Data Classes | Maturity | Description |
| --- | --- | --- |
| **Constraints** - rules or set of rules that must be satisfied for the modeled Categorical Variant |  |  |
| Defining Allele | Trial use | Specific VRS Alleles |
| Defining Location | Trial use | Sequence Locations |
| Copy Count | Trial use | Exact or range of copies |
| Copy Change | Draft | Relative assessment of copies |
| Feature Context | Draft | General concepts (e.g., genes, protein markers) |
| Function | In progress | Protein functional consequence |
| Adjacency | In progress | Adjoining of two sequences |
| **Categorical Variant Types** - schemas for representing types of variation using Constraints |  |  |
| Canonical Allele | Trial use | Genomic DNA changes |
| Protein Sequence Consequence | Trial use | Amino acid changes |
| Categorical CNV | Draft | Copy number changes |
| Function Variant | In progress | Gain / Loss of function variants |
| Gene Fusion | In progress | Adjoining of two genes |

This constraint-based model directly addresses the limitations of earlier systems. Unlike top-down typologies, Cat-VRS is compositional and extensible: new categories can be formed by combining existing constraint types without requiring schema changes. Unlike purely molecular representations, Cat-VRS accommodates variant classes that are not sequence-resolvable, including functional categories, structural variant families, and clinically defined biomarker groups. Most importantly, constraints are fully computable objects, enabling entailment and matching operations that allow programmatic determination of when an observed variant satisfies a categorical definition, even when the category’s label is ambiguous or inconsistently defined across resources.

## Discussion

The absence of a unified representational model for categorical variation has directly contributed to the variant interpretation bottleneck in precision medicine by limiting search, consistent reasoning, and scalable knowledge integration^3,16,28,49^. Historically, queries for clinically relevant categories of variation have relied on ad hoc text matching, manually curated lists, or knowledge base-specific definitions that cannot be generalized across systems^16,24,46^. For example, “activating *BRAF* variants”, “any *PTEN* loss-of-function variant”, or “variants affecting *EGFR* exon 19”. These limitations impact variant interpretation and thus treatment eligibility determination, clinical trial enrollment, and functional evidence assessment, and furthermore complicate audibility and validation of clinical decision support (CDS) pipelines when category definitions are implicit or inconsistently encoded. Categorical variant representation of individual sequence variants has remained problematic as well due to the multiplicity of possible expressions for a single alteration. Cat-VRS builds upon VRS to retain the benefits of precise, unambiguous representation of individual variants while also modeling categorical variants as sets of discrete properties through constraints that filter the space of genomic alterations. In doing so, Cat-VRS provides the semantic layer necessary to harmonize category definitions across genomic knowledge resources.

Cat-VRS closes longstanding interoperability gaps in precision medicine across several application domains. In category-based variant querying, the formalization of categorical variants as constraint-defined domains enables the programmatic evaluation of categorical membership across nomenclature systems, transcripts, and structural encodings, while also establishing a foundation for integration into variant annotation pipelines and analytic workflows. In clinical trial matching, eligibility criteria based on categorical descriptions that span biological levels can be encoded as explicit constraint-based definitions, allowing eligibility rules to be evaluated directly against assayed variants and facilitating interoperability with standards such as HL7 FHIR. In clinical decision support, Cat-VRS enables programmatic entailment testing between observed variants and categorical concepts used in therapeutic guidelines and genomic knowledge statements. Fundamentally, Cat-VRS’ capabilities support interoperable, dynamically updated evidence retrieval for categorical variants within a standards-based infrastructure^12,50^.

In practice, the integration of constraint-based modeling and VRS identifiers substantially reduces the burden of variant knowledge curation and maintenance. Curators may thus focus on assessing evidence quality and clinical significance rather than on representational alignment, yielding clearer provenance trails that support validation and regulatory review of clinical decision support pipelines.

The v1.0.0 release of Cat-VRS^51^, published in June 2025, provides the first stable set of constraint classes for representing categorical variants by a specific allele, sequence location, and copy number, alongside reference examples for how to represent canonical alleles and protein sequence consequences using the specification (Table 1). Several additional constraint classes are actively being prepared for release: the representation of categorical copy-number, functional impact, definition by concepts (for example, a specific gene), adjacency for representing structural events such as fusions and rearrangements, and the representation of pathogenicity and oncogenicity variant categories. Together, these constraints expand Cat-VRS toward fully capturing the breadth of categories used across genomics knowledgebases, variant interpretation workflows, and clinical reporting. Support for complex categorical variants, where multiple categorical variants are associated by a boolean operator, is a subsequent focus and will enable richer representational capabilities, such as enabling the representation of categories such as “BRAF V600 variants” (any residue at position 600 except valine).

Cat-VRS has been, and continues to be, developed as an open community standard grounded in implementer-driven development. Since October 2023, the working group has held more than 45 open development meetings with participation from over 57 individuals across 37 institutions, representing 9 countries across 5 continents. This diverse engagement has been essential for identifying use case requirements, developing product features, and guiding design decisions. Importantly, continued maturation of the specification, both in the expansion of new constraint types and in the progression of existing ones from draft to trial use to normative maturity, relies on broad adoption. Under GA4GH governance for genomic knowledge standards, product features advance in maturity status only when they are used, evaluated, and endorsed by real-world users (currently represented by 12 registered implementations). As more organizations adopt Cat-VRS, they both benefit from and contribute to this maturation cycle by stress-testing the model, uncovering edge cases, shaping new feature requests, and participating directly in the collaborative development process.

By providing a principled, computable representation of categorical variation, Cat-VRS offers a foundation for interoperability across genomic knowledgebases, variant interpretation pipelines, clinical trial infrastructures, and decision-support systems. Building on our survey of categorical variants across the biomedical ecosystem, our formal analysis of CIViC variant types, and the design and implementation of a novel constraint-based model, we have demonstrated how Cat-VRS enables computational capabilities that were previously inaccessible, from category-based search to knowledgebase harmonization to programmatic trial matching and clinical decision support. Future efforts will focus on accelerating adoption, enhancing implementation guidance, and deepening integration with other GA4GH products such as Beacon. Ultimately, Cat-VRS seeks to reduce the interpretation bottleneck in precision medicine by enabling reliable, computable connections between observed variants and the clinically meaningful categories to which they belong. We invite the broader community to join us in implementing, testing, and refining this open standard, and to contribute to a scalable, interoperable future for genomic knowledge.

## Acknowledgements

We acknowledge Beatrice Amos (Global Alliance for Genomics and Health) for her supervisory and project administration support to the Cat-VRS product development group from its inception as study group through v1.0 public release of the specification and the production of the present manuscript. We also acknowledge Salem Bajjali (Mayo Clinic) for his role in reviewing and editing the manuscript.

Furthermore, we acknowledge the vital roles of Sue Mockus (Baylor Genetics), Michael Baudis (University of Zurich), and Tristan Nelson (Geisinger), who served as the product review committee within GA4GH to oversee and guide the development of Cat-VRS the original landscape analysis through product development and eventual public 1.0 release.

Research reported in this publication was supported by the National Institutes of Health [R01 EB30529, R35 HG011899, R35 HG011949, U24 CA275783, U24 CA305456, U24 HG011025], the Wellcome Trust [220544/Z/20/Z], Medical Research Council [MC_PC_19024], National Institute of Health Research, Canadian Institute for Health Research [202506NDG

CIHR-IRSC:0550017031], and the Six Four Foundation.

GA4GH is supported by the National Institutes of Health [U24 HG011025], the Wellcome Trust [220544/Z/20/Z], Medical Research Council [MC_PC_19024], National Institute of Health Research, and Canadian Institute for Health Research [202506NDG CIHR-IRSC:0550017031].

## Ethics declarations

B.R. has institutional patents filed on methods for clinical interpretation. R.D.is employed by Elimu Informatics. C.P. is employed by Epic Systems. B.P. serves as a volunteer on the Qiagen Somatic Advisory Board. P.K. is employed by Velsera. A.R. is employed by OakBioinformatics LLC. E.M.V.A. serves as a consultant or on scientific advisory boards of Novartis Institute for Biomedical Research, Serinus Bio, TracerBio, Cellyrix. E.M.V.A. has received research support (to institution) from Novartis and BMS. E.M.V.A. has equity in Tango Therapeutics, Genome Medical, Genomic Life, Enara Bio, Manifold Bio, Microsoft, Monte Rosa, Riva Therapeutics, Serious Bio, Synapse, TraceBio, and Cellyrix. E.M.V.A. has institutional patents filed on chromatin mutations and immunotherapy response, and methods for clinical interpretation and performs intermittent legal consulting on patents for Foaley & Hoag. The remaining Authors declare no competing interests.

## Online Methods

### Landscape survey of categorical variant usage

A survey of genomic knowledge resources that utilize categorical variants was undertaken in September 2023 - January 2024 as part of the GA4GH Categorical Variation Study Group prior to Cat-VRS development. A list of 34 resources were contributed to by study group participants during this period, representing input from 25 individuals across 16 institutions. Each resource was categorized as either an API (2 of 34 resources), application or knowledgebase (20), data model (6), initiative (1), terminology or identifier system (3), or other (2). Resources were further classified as either according to whether they stored categorical variants (17 of 34 sources), worked with them (17), or neither (1).

### Categorical variants associated with US FDA oncology regulatory approvals

Oncology approvals from the US Food and Drug Administration involving biomarkers were downloaded from the Molecular Oncology Almanac’s June 12th, 2025 release^46^. A total of 237 approvals were downloaded and reformatted from JSON into a tab delimited file, containing the drug’s application number, drug brand name, drug generic name, package insert URL, and regulatory approval (‘indication’) text. Each approval was manually reviewed and labeled with either TRUE or FALSE if the indication involved a set of genomic variations or not. The categorical variant(s) involved were also annotated, delimited by semi-colons if multiple categorical variants were associated with a single approval.

A total of 44 distinct categorical variants were identified, spanning categories of specificity involving: allele origin, resulting amino acid, clinical impact, exon location, fusion partner, involved gene, involved gene set, functional impact (pathogenicity), and variant type. A JSON file was created with a record for each categorical variant, containing keys for the relevant categories, relevant approvals, and count of relevant approvals.

The co-mut package was then used to create a heatmap of all variants with involved categories, paired with a horizontal bar chart for the number of relevant approvals per categorical variant (Fig 2a, Crowdis et al. 2020).

### Multiplicity of Alleles in ClinVar

The hgvs4variation.txt tab delimited output was downloaded from ClinVar’s FTP site (https://ftp.ncbi.nlm.nih.gov/pub/clinvar/tab_delimited/); specifically, the 2024-03-31 release, which was last modified on 2025-06-30 (md5: d680fb3ffbee054a601785ba905c6a33). A histogram of the total HGVS expressions (both nucleotide and protein) per Allele ID was generated. The x-axis was adjusted to use log-spaced bins for interpretability (Fig. 2b).

## Data and code availability

This study did not involve the generation of biological or chemical data.

The specification described by this study is available online at https://cat-vrs.ga4gh.org, which is generated from the specification codebase at https://github.com/ga4gh/cat-vrs. The v1.0.0 specification has been deposited at Zenodo, at https://doi.org/10.5281/zenodo.17093422. All code and generated display items used in the present study were completed using Python 3.12 and are openly available through this study’s public GitHub repository, at https://github.com/ga4gh/2025-catvar-call-to-action. Both the Categorical Variant Representation Specification and this study’s GitHub repository are distributed under the Apache-2.0 license.

## Display Items

## Supplementary Information

**Supp. Fig. 1:**
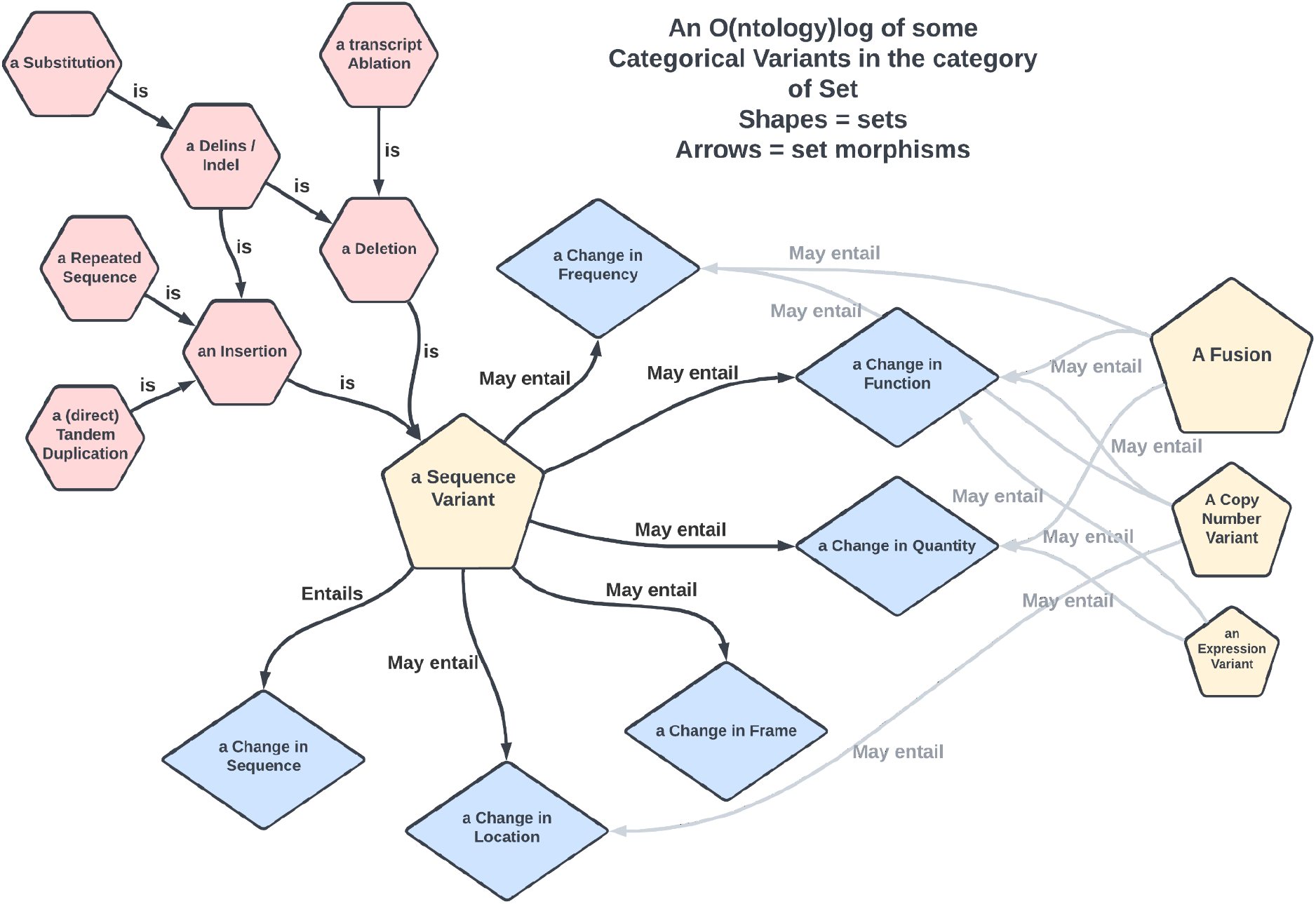
Olog representation of CIViC categorical variant types. This ontology log (olog) depicts a subset of CIViC’s categorical variant types as set objects, with arrows representing definitional relationships (set morphisms) among them. By formalizing these categories and the properties they entail, the olog clarifies how CIViC’s variant types relate to one another and salient semantic properties. This structural analysis provided the basis for identifying the minimal properties that govern category membership and informed the development of the typology and constraint-based representation used in Cat-VRS.

## Supplementary Table 1

**Supplementary Table 1: Landscape of resources relevant to categorical variation**. A summary of genomic knowledgebases, clinical resources, and data models that store or use categorical variants, or that would benefit from interoperability with Cat-VRS.

